# The Social Role of Self-Control

**DOI:** 10.1101/2020.10.26.354852

**Authors:** R.I.M. Dunbar, Susanne Shultz

## Abstract

The capacity to inhibit prepotent actions (inhibitory self-control) plays an important role in many aspects of the behaviour of birds and mammals. Although a number of studies have used it as an index of foraging skills, inhibition is also crucial for maintaining the temporal and spatial coherence of bonded social groups. Using three different sets of comparative data, we show that, across primate species, the capacity for self-control correlates better with the demands of social contexts than with the demands of foraging contexts, whereas a more generalised capacity for causal reasoning correlates better with foraging contexts. In addition, we confirm the Passingham-Wise Conjecture that the capacity for self-control is unique to anthropoid primates. These results suggest that the capacity for self-control most likely evolved because it was crucial for the evolution of bonded social groups.

**Significance Statement:** The capacity for self-control has commonly been viewed as an index of foraging skills. In fact, it plays a much more important role in the social domain by enabling groups of animals to maintain social cohesion as they travel through time and space. In this respect, it is particularly important for species that live in stable bonded social groups (congregations). We show that, in this respect, it is uniquely characteristic of the anthropoid primates, in contrast to other kinds of reasoning tasks such as causal reasoning on which primates often perform no better than other birds and mammals.

## Introduction

Anthropoid primates have evolved a form of bonded sociality that allows them to live in large, stable, cohesive social groups of a kind that is rare among birds and mammals, being otherwise found only in a small number of (mainly species-poor) orders such as equids, elephantids, tylopods and delphinids (Shultz & Dunbar 2007, 2010; Sutcliffe et al. 2016; Dunbar & Shultz 2021a). Bonded social groups face two significant challenges in terms of maintaining social cohesion in the face of the forces that normally cause herds and flocks to fragment and disperse. One is the need to manage the stresses experienced when living in close physical proximity with others that would otherwise cause individuals to leave and find smaller groups (Dunbar & Shultz 2021b). The other is the coordination problem that arises when failure to synchronise activity schedules during foraging results in groups becoming dispersed and eventually fragmenting.

In species that form aggregations (unstable flocks or herds), differences in the rate of gut fill inevitably result in animals’ time budgets becoming desynchronised when some individuals want to rest whereas others want to continue feeding (Conradt 1998; Conradt & Roper 2000; Dunbar & Shi 2008; Conradt et al. 2009). More generally, individual differences in foraging strategy or food preferences may result in some individuals wanting to move to another food patch while others prefer to stay where they are (Krause & Ruxton 2002). When the bonds between group-members are weak, animals will simply drift apart and groups will fragment (Ruckstuhl & Kokko 2002; Ruckstuhl & Neuhaus 2002; Calhim et al. 2006; Dunbar & Shi 2008). To be able to live in stable groups, animals need to be able to resist the temptation to continue feeding when the rest of the group wants to go to rest, or to rest when others want to continue foraging (King & Seuer 2009, 2011; Strandburg-Peshkin et al. 2015, 2017; Aguilar-Melo et al. 2020; Papageorgiou & Farine 2020; Harel et al. 2021; Schweinfurth et al. 2022; Schaposnik et al. 2023). In this context, the capacity to inhibit prepotent responses (self-control) is likely to play a crucial role in allowing individuals to resist the temptation to carry on with their currently preferred activity in order to remain synchronised with their core social partners.

Many studies have, however, emphasised a role for self-control in foraging (MacLean et al. 2014; Stevens 2014; Rosati 2017) on the grounds that, in order to forage optimally, animals have to be able to forego a less valuable immediate reward in order to gain a more desirable future one (Stephens & Krebs 1986). By implication, the capacity for self-control, necessarily dependent on a large brain, evolved to allow primates (in particular) to engage in more complex foraging strategies. A number of studies have, however, questioned whether the capacity to inhibit prepotent responses requires large primate-sized brains (see Kabadayi et al. 2016; Jelbert et al. 2016; van Horick et al. 2018; Isakson et al. 2018). These studies have claimed that self-control is a widely distributed cognitive skill, and not particularly dependent on having a large brain. The suggestion that birds may be able to engage in self-control is, however, at odds with the claim that the capacity to inhibit behavioural responses (self-control) may in fact be a uniquely anthropoid primate trait that depends on a brain region (the frontal pole, or Brodman areas BA9/10) that is only found in this taxon (Passingham & Wise 2012). We refer to this as the Passingham-Wise Conjecture.

Although the Passingham-Wise Conjecture has yet to be tested on a broad taxonomic scale, neuropsychological experiments with both monkeys and humans suggest that foraging decisions that involve a change in behavioural strategy in response to a change in the reward schedule typically activate the ACC (anterior cingulate cortex, a brain region associated with error detection and identifying violations of expectation: Kennerley et al. 2006; Holroyd et al. 2009), whereas neuroimaging and lesion studies in humans and primates suggest that inhibition depends on the frontal pole, in particular the ventrolateral prefrontal cortex (Aron et al. 2004; Passingham & Wise 2012; Passingham 2021). This implies that foraging decisions may not necessarily involve the capacity for self-control, in turn implying that this cognitive ability may have evolved for other purposes.

We here test two hypotheses: (1) that, in primates, the capacity for behavioural self-control is more strongly correlated with the demands of maintaining social cohesion than with foraging decisions and (2) that the capacity for self-control is uniquely characteristic of the anthropoid primates. To test the first hypothesis, we analyse comparative experimental data on several well established cognitive tasks for a range of primate species using three different datasets (Amici et al. 2008; MacLean et al. 2014; Stevens 2014). Although these various tasks correlate reasonably well with each other, doubts have been expressed as to whether they actually index the same underlying cognitive mechanism. This has been discussed in particular with respect to the ‘cylinder task’ (Kabadayi et al. 2017), a form of detour task that has been widely used as an inhibition task (MacLean et al. 2014; Kabadayi et al. 2016; Jelbert et al. 2016; van Horick et al. 2018; Isakson et al. 2018). The cylinder task asks animals to choose between two ends of a tube in order to access a food reward, one of which is more accessible than the other (either because one end is blocked or because the reward is nearer one end). Consideration of the task demands involved suggests that it might, in fact, be better characterized as a causal reasoning task. Because of these doubts, we first determine whether the different tasks that have been used in these studies measure the same underlying cognitive ability. For this, we use the dataset provided by Amici et al. (2008) since, although they include fewer species, they provide data on five different tasks, all of which they refer to as ‘inhibition tasks’. We show that, in fact, they form two distinct clusters: a set of tasks that do seem to index self-control and a set of tasks that are analogous in their design to the cylinder task and are best described as indexing causal reasoning. We therefore treat these two classes of tasks separately.

We then determine, for each of the three datasets separately, whether these cognitive variables cluster better with a pair of variables that reflect the demands of maintaining group coherence during foraging (group size and day journey length) or with a pair of variables that reflect the demands of foraging decisions (diet and home range size) (for details, see *Methods*). Having established which variables cluster with which, we then use mediation analysis to determine the likely causality involved. Finally, we use the full MacLean et al. (2014) dataset (which includes data for a wide range of bird and mammal species in addition to primates) to test the Passingham-Wise Conjecture that only anthropoid primates have the capacity to inhibit prepotent responses, whereas we expect detour tasks that index causal reasoning to be widely characteristic of other mammals and birds primates as well as primates.

## Methods

### Data

We use data from three independent studies (Amici et al. 2008; MacLean et al. 2014; Stevens 2014) that provide experimental data on the capacity to inhibit prepotent responses based on different behavioural tasks for between seven and 18 primate species (Table 1). Amici et al. (2008) and MacLean et al. (2014) carried out a series of experiments on different species using the same experimental design; Stevens (2014) collated data from the literature where different studies had used a similar task (a Go/No-go task) on a variety of different taxa. Since they involve different tasks using different metrics, we analyse the three datasets separately. We sourced mean group size for species from Dunbar et al. (2018), percentage of fruit in the diet (except for *Macaca mulatta*: see Table S3) from Powell et al. (2017) and day journey length (in km) and home range size (in ha) from Smuts et al. (1987) and Campbell et al. (2007) and primary sources therein (for exceptions, see *ESM*).

**Table 1.**
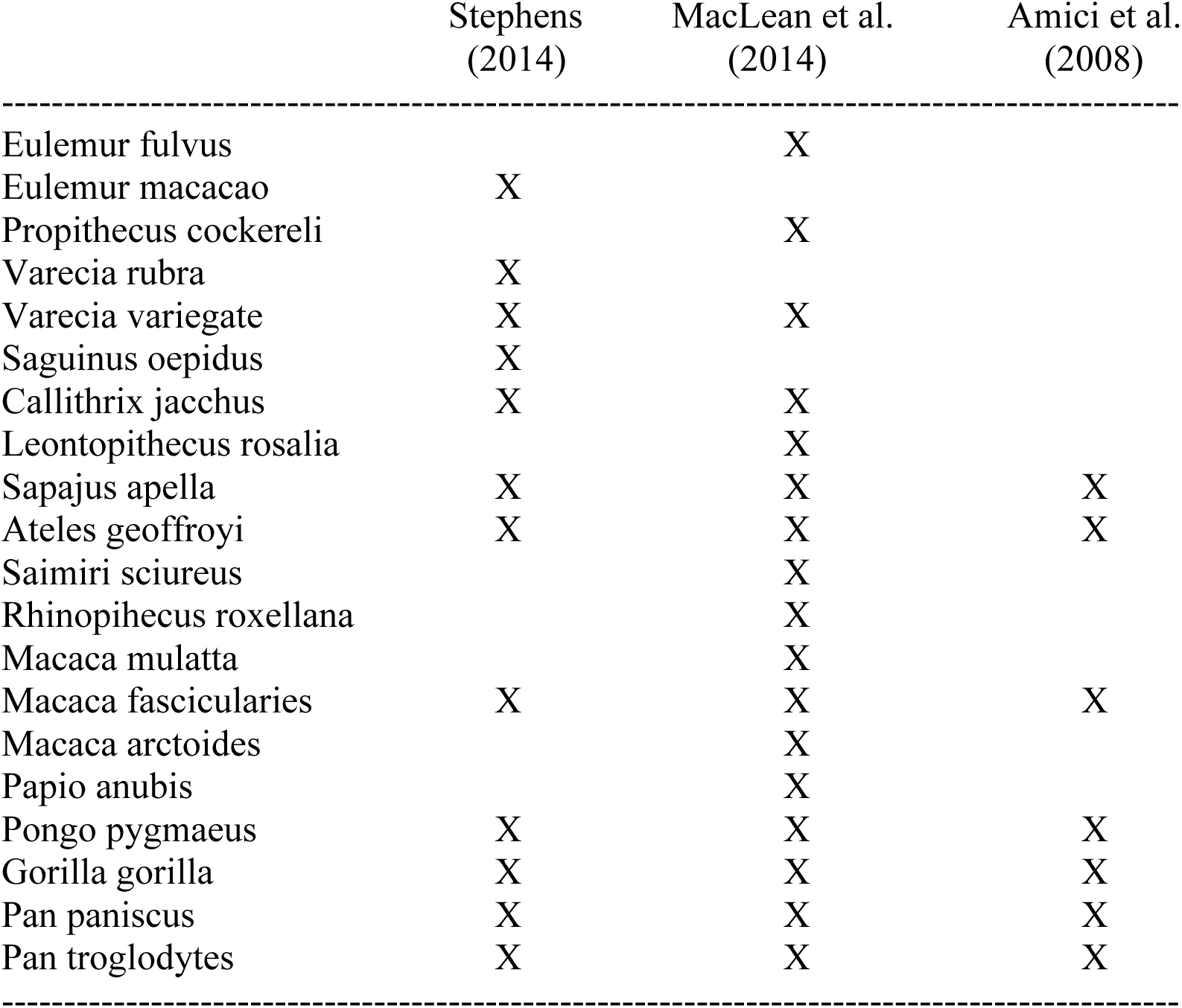
Primate species sampled.

**Table 2.** Regression analysis of fission index for 25 different *Papio* study sites.

| Variable | slope | standardised $\beta$ | df | t | p |
| --- | --- | --- | --- | --- | --- |
| Overall model: | $r^2=0.664$, $F_{3,22}=14.47$, $p<0.0001$ | | | | |
| SD(group size ) | 0.481 | 0.546 | 22 | 2.20 | 0.039 |
| SD(day journey) | 0.373 | 0.423 | 22 | 2.17 | 0.041 |
| Interaction | -0.182 | -0.207 | 22 | -1.07 | 0.295 |

We exclude *Daubentonia* and *Papio hamadryas* from the MacLean et al. dataset because of uncertainties about their correct group sizes. For detailed discussion on this, and on group size for *Pongo*, see *ESM*.

### The data are given in *ESM DATASET-1*

As indices of foraging demand for the main analysis, we use the percentage of fruit in the diet and the size of the home range (or territory), both of which have frequently been used to test foraging demand hypotheses (Clutton-Brock & Harvey 1980; Dunbar 1992; MacLean et al. 2014; Stevens 2014; Powell et al. 2017). Fruits are much less predictable than foliage, and are usually viewed as representing a more cognitively challenging diet than leaves (Clutton-Brock & Harvey 1980). Similarly, large home ranges are assumed to be cognitively demanding both in terms of the mental mapping skills required to devise an optimal foraging pathway between food sites and in the fact that foraging animals have to choose between near and distant locations on the basis of nutrient value (MacLean et al. 2014; Stevens 2014). In primates, both percent fruit in the diet and range size are strongly influenced by habitat conditions and hence impact on nutrient acquisition (Hill & Dunbar 2002; Lehmann et al. 2008; Dunbar et al. 2009, 2019; Ménard et al. 2013). If inhibition relates to foraging efficiency, performance on such tasks should correlate positively with these ecological indices: the more patchy and dispersed the distribution of food resources, the more animals will need to be able to inhibit the temptation to stay in the current resource patch rather than delay the rewards from going to a richer one that is further away.

In respect of the social domain, we focus on the role that self-control is likely to play in ensuring group cohesion during foraging. For social species, the larger the group and the longer the day journey the more important self-control will be for maintaining group cohesion. Coordination problems increase as a function of both group size and the distance animals have to travel since both make it more likely that individuals’ activity cycles will progressively drift out of synchrony (Conradt & Roper 2000; King & Cowlishaw 2009). Baboons provide a well studied example: the risk of group fragmentation will increase as day journey lengths get longer and group sizes get larger (Fig. 1; Tables S1-S2). The fissioning index correlates significantly with both group size (τ=0.642, N=26, p<0.001) and day journey length (Kendall’s τ=0.655, N=26, p<0.001). To determine whether there is an interaction effect between the two independent variables, we transformed group size and day journey length to standard normal deviates, and ran a multiple regression with an interaction effect. The results are given in Table 1. There are significant main effects, of approximately equal weight, but no interaction effect. If inhibition is primarily a social skill that influences group cohesion, it should correlate positively with these indices. We would not expect it to correlate with indices of foraging.

**Fig. 1.**
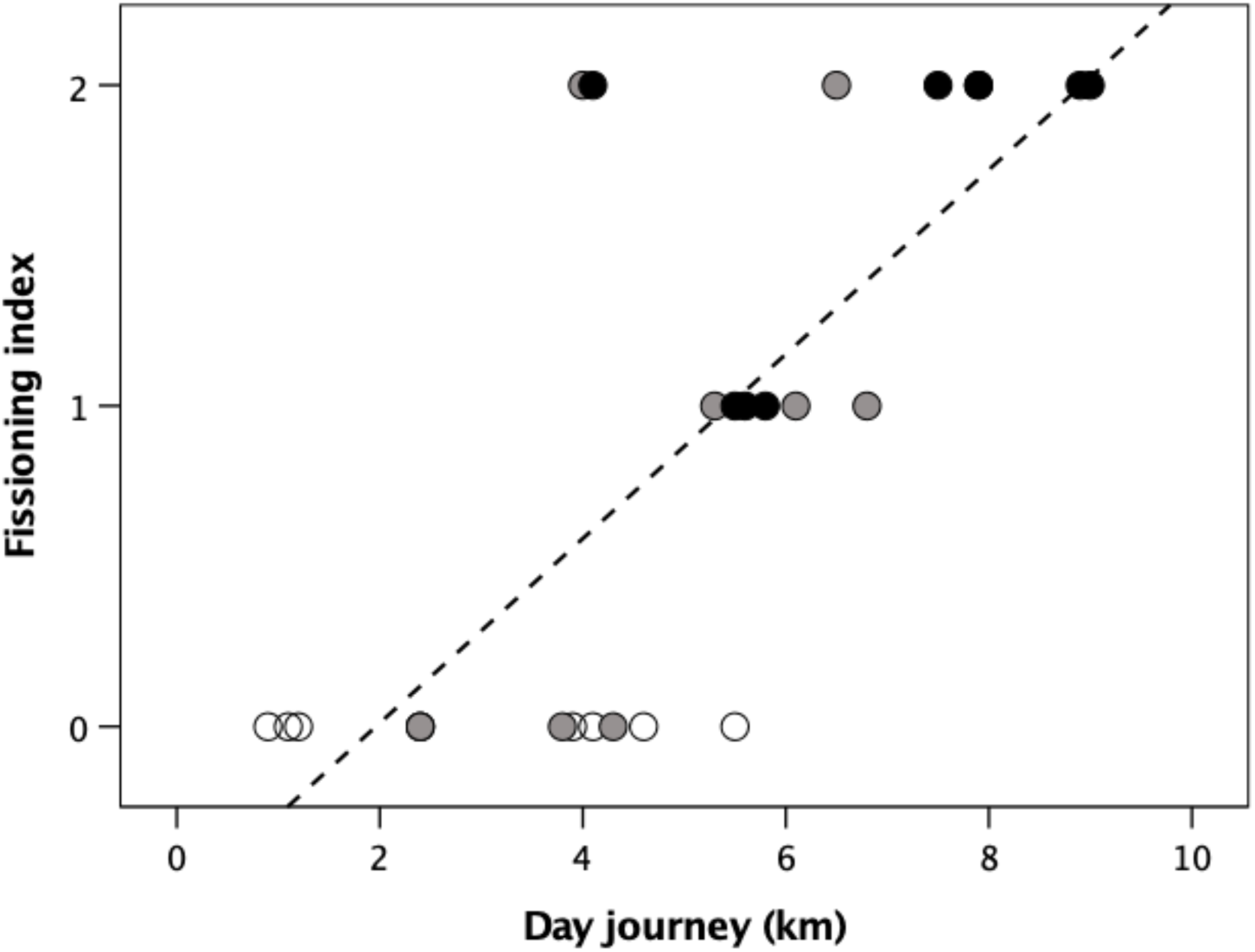
Fissioning index for individual *Papio* baboon populations as a function of day journey length. Unfilled symbols: group size <35; grey symbols: group size 35-75; filled symbols: group size >75. For definition of fissioning index, see Table S2. The data are given in Table S1.

It is important to be clear about the difference between range size and day journey length here since, viewed superficially, both look like foraging-relevant variables. In fact, functionally they are very different, and especially so for primates. Unlike herding species, most primates are frugivores: they do not forage semi-randomly in their environment in the way grazers do, but rather move from one discrete resource patch to another, often at some considerable distance (Altmann & Altmann 1970; Sigg & Stolba 1981). Range size determines the number of patches available to a group, but does not, of itself, determine the number of patches visited each day or the length of the day journey. Day journey length, by contrast, is largely a consequence of the size of the group and the number of patches the group has to visit to satisfy its collective nutritional requirements. In other words, range size defines the distribution of food sources that animals can choose between and hence the choices they make on where to forage, whereas day journey length is simply the means for visiting the required number of patches each day (but *not* which patches to visit). The first is a resource choice issue, the second a routing issue. Only the second has significant implications for maintaining group cohesion.

To test the Passingham-Wise conjecture, we use the data given by MacLean et al. (2014) on the A-not-B and cylinder tasks for a wide range of 36 mammalian and avian species. In this case, we ask simply whether major taxonomic groups differ in their performance on the A-not-B (inhibition) task or the cylinder (causal reasoning) task. The data are given in *ESM DATASET-2*.

### Statistical analysis

Although multiple regression might seem the obvious method for testing hypotheses of this kind, the format of the standard regression model would oblige us to regress the cognitive cause (inhibition skill) on the four ecological and social outcome variables just as MacLean et al. (2014) and Stevens (2014) in fact do. However, doing so reverses the natural causality (i.e. by assuming that behaviour determines cognitive ability rather than cognitive capacity determining, or constraining, behaviour). Because this asks a very different question, it can yield very different results that are susceptible to misinterpretation (Dunbar & Shultz 2023). A statistically more elegant approach is to use principal components analysis (PCA) to ascertain which variables covary (i.e. cluster together). PCA avoids the need to presumptively specify the causal relationships between variables (Mackinnon et al. 2007).

We do not correct for phylogeny in these analyses because reliable methods for doing so with PCA do not yet exist. In any case, phylogenetic correction should only be used when there is reason to believe that phylogenetic inertia might artificially inflate the degrees of freedom in a statistical analysis. Adding additional, causally irrelevant variables to a statistical analysis without justification breaches the demand for parsimony and reduces statistical power unnecessarily. In the present case, previous conventional regression analyses of both the behavioural and ecological variables (Shultz & Dunbar 2007, 2010) and the cognitive variables (Stevens 2014; Amici et al. 2008) have found no effect of phylogeny on the results. This is because, in primates, the phylogenetic signals for group size, percent fruit in diet, home range size and day journey length are all close to zero (Kamilar & Cooper 2013). This largely reflects the fact that primates are behaviourally extremely flexible (Strier et al. 2014), and in consequence most of the variance in their behaviour (including group size) is environmentally, and not genetically, determined (Dunbar et al. 2009; Dunbar & Shultz 2021b).

All analyses were undertaken in SPSS v.29.

## Results

We first use PCA to determine whether Amici et al.’s five cognitive tasks index the same underlying variable. Table 3 gives the results. It is clear that the five tasks partition into two subsets. One subset includes the A-not-B task (the same task as used by MacLean et al. 2014) and a delayed gratification task (another well known inhibition task). The other subset includes their Middle cup, Plexiglass holes and Swing door tasks. Although these latter tests are presented as assays for inhibition (and were so presented by the sources from which the designs were taken), in fact they are best understood as tests of causal understanding. Understanding that pushing a door will knock the reward off the shelf behind whereas pulling the door will allow the animal to reach through and get the reward, for example, has much more to do with understanding causality than being a matter of inhibition. Since each of these two subsets correlates well internally, we averaged the values for the respective subsets, referring to the first subset as Inhibition tasks and to the second subset as Causal reasoning tasks. We compared these two indices with the same species’ performances on the social inhibition task of Amici et al. (2018) and the A-not-B and cylinder tasks used by MacLean et al. (2014). Except the Amici social inhibition and causal reasoning tasks, none of these tasks correlate significantly with each other, and several correlate negatively (Table S4). Thus, despite having been presented in different publications as indices of inhibition, it seems that they in fact probably tap into very different cognitive skills. We therefore treat them separately. To test whether the various cognitive variables are associated with any of the four behavioural ecology variables, we ran a separate PCA for each cognitive task. (Bivariate correlations between the Stevens and MacLean cognitive tasks and the four ecological outcome measures for the two larger datasets are given in Table S5 and Fig. S1.) For the Amici et al. (2008) dataset, we included both cognitive indices in the same analysis; the analysis for the inhibition task on its own is given in Table S6. With eigenvalues set to λ>1, two factors are extracted for all five datasets, explaining >70% of the variance in each case (Table 4). The three self-control (inhibition) tasks, group size and day journey length consistently load together on the same factor with high loadings, while diet is placed either in a separate factor or with the cylinder task. When the A-not-B and cylinder tasks are both included in the same analysis, the cylinder task loads more strongly with diet. Similarly, when the Amici et al. (2008) inhibition and causal reasoning tasks are included in the same analysis, home range and causal reasoning ability are placed together in a separate factor, while diet loads on its own. In general, range size typically loads weakly across factors.

**Table 3.** Factor loadings (with varimax rotation) for the five cognitive tasks in the Amici et al. (2008) dataset.

| 730 | Tasks | 1 | Factors<br>2 |
| --- | --- | --- | --- |
|  | A-not-B | **0.925** | -0.052 |
|  | Middle cup | 0.224 | **0.763** |
| 735 | Plexiglass hole | 0.472 | **0.731** |
|  | Swing door | -0.193 | **0.833** |
|  | Delayed gratification | **0.898** | 0.264 |
|  |  |  |  |
|  | Species sampled | 7 |  |
| 740 | Variance explained | 77.1% |  |

Extraction based on $\lambda = 1.0$ . Bold font indicates variables that have a strong positive load on the same factor.

**Table 4.** PCA factor loadings (with varimax rotation) for the variables for each of the three datasets.

|  |  | Stevens (2014) |  | MacLean et al. (2014) |  |  |  | Amici et al. (2008) |  |  |
| --- | --- | --- | --- | --- | --- | --- | --- | --- | --- | --- |
|  |  | Go/No-go task |  | A-not-B task only |  | A-not-B and cylinder tasks |  | Inhibition† and Causality tasks‡ |  |  |
| 750 | Factors: | 1 | 2 | 1 | 2 | 1 | 2 | 1 | 2 | 3 |
| ----- |  |  |  |  |  |  |  |  |  |  |
| 755 | Inhibition task | **0.898** | 0.249 | **0.855** | 0.053 | **0.849** | 0.189 | **0.871** | 0.308 | -0.122 |
|  | Cylinder task |  |  |  |  | 0.593 | **0.740** |  |  |  |
|  | Causal reasoning tasks† |  |  |  |  |  |  | 0.191 | **0.955** | 0.010 |
|  | Group size | **0.885** | 0.102 | **0.900** | 0.031 | **0.888** | 0.030 | **0.915** | 0.027 | 0.250 |
|  | Day journey | **0.969** | -0.090 | **0.895** | -0.180 | **0.915** | -0.091 | **0.915** | -0.237 | -0.238 |
| 760 | Diet (% fruit) | -0.018 | **0.778** | 0.020 | **0.971** | -0.156 | **0.931** | -0.036 | 0.102 | **0.984** |
|  | Range size | 0.148 | **0.714** | 0.498 | 0.208 | 0.485 | 0.356 | -0.123 | **0.941** | 0.145 |
|  | Species sampled | 13 |  | 18 |  | 14 |  | 7 |  |  |
|  | Variance explained | 74.9% |  | 72.3% |  | 75.7% |  | 92.8% |  |  |

Extraction based on $\lambda = 1.0$ . Bold font indicates variables that have a strong positive load ( $>0.700$ ) on the same factor.
† mean of two inhibition tasks (A-not-B and delayed gratification tasks)
‡ mean of three detour tasks (middle cup, plexiglass and swing door)

In case our choice of group size for *Pongo* distorted the results, we re-ran the three PCAs excluding this genus. Table S7a indicates that the results do not change (other than moving range area into the inhibition factor and leaving diet isolated on the second factor). Table 7b confirms that including *Papio hamadryas* in the MacLean et al. analysis with alternative grouping sizes does not change the main results. Notice that, although the overall fit is slightly lower, the loadings for the smaller group size are a very close match to those in Table 4. *P. hamadryas* was not included in the other two datasets, so this species could not have biased the results in either of these cases. Table S8 confirms that these results hold when the data are analysed as genus level means, indicating a very limited influence for phylogeny. In sum, it seems that potential confounds have not distorted the results.

Amici et al. (2018) provide data on an explicitly social inhibition task (will animals resist taking a reward if they are in the presence of a more dominant individual?) for six of the same animal populations tested in Amici et al. (2008). We ran a PCA for this index on its own with the four ecological variables, and in combination with Amici et al.’s (2008) inhibition and causal reasoning indices (Table 5). While the inhibition index loads once again with group size and day journey length as in Tables 4 and S6, the social inhibition task loads on the same factor as causal reasoning and diet (with home range being again somewhat ambivalent in its loadings), suggesting that it may not be indexing the same underlying phenomenon as the A-not-B and Go/No-go tasks; instead, it may in fact be indexing something closer to causal reasoning (in effect: the dominant animal will attack me if I take the food).

**Table 5.** PCA factor loadings (with varimax rotation) for the three cognitive variables from Amici et al. (2008) and Amici et al. (2018)

|  | Factors: | Social inhibition only |  | All three cognition tasks |  |
| --- | --- | --- | --- | --- | --- |
|  |  | 1 | 2 | 1 | 2 |
| 780 | Social inhibition task* | **0.880** | 0.297 | **0.855** | 0.371 |
|  | Inhibition tasks † |  |  | 0.234 | **0.922** |
|  | Causal reasoning tasks ‡ |  |  | **0.970** | -0.055 |
|  | Group size | 0.211 | **0.919** | 0.154 | **0.899** |
|  | Day journey | -0.213 | **0.955** | -0.268 | **0.930** |
| 785 | Diet (% fruit) | **0.880** | -0.237 | **0.834** | 0.080 |
|  | Range size | **0.838** | -0.010 | -0.217 | **0.893** |
|  | Species sampled | 6 |  | 6 |  |
|  | Variance explained (%) | 84.9 |  | 86.2 |  |

Extraction based on $\lambda = 1.0$ . Bold font indicates variables that have a strong positive load on the same factor.
\* from Amici et al. (2018)
† mean of two inhibition tasks (A-not-B and delayed gratification tasks); from Amici et al. (2008)
‡ mean of three causal reasoning tasks (Middle cup, Plexiglass hole and Swing door); from Amici et al. (2008)

To evaluate the relationship between self-control and its associated social variables in more detail, we ran a mediation analysis with the inhibition index with the largest sample (the MacLean et al. A-not-B task) as the independent variable and group size and day journey length alternately as dependent variable and mediator. A Sobel test indicates that a model with group size as mediator and day journey as dependent variable (Fig. 2) yields a significant model (z=2.893, p=0.0038). This model is significantly better than one with group size as the dependent variable and day journey as the mediator (z=0.022, p=0.982) or one with inhibition as the dependent variable (z=0.070, p=0.944). This suggests that the capacity to inhibit behaviour (self-control) determines group size, and group size then determines day journey length.

**Fig. 2.**
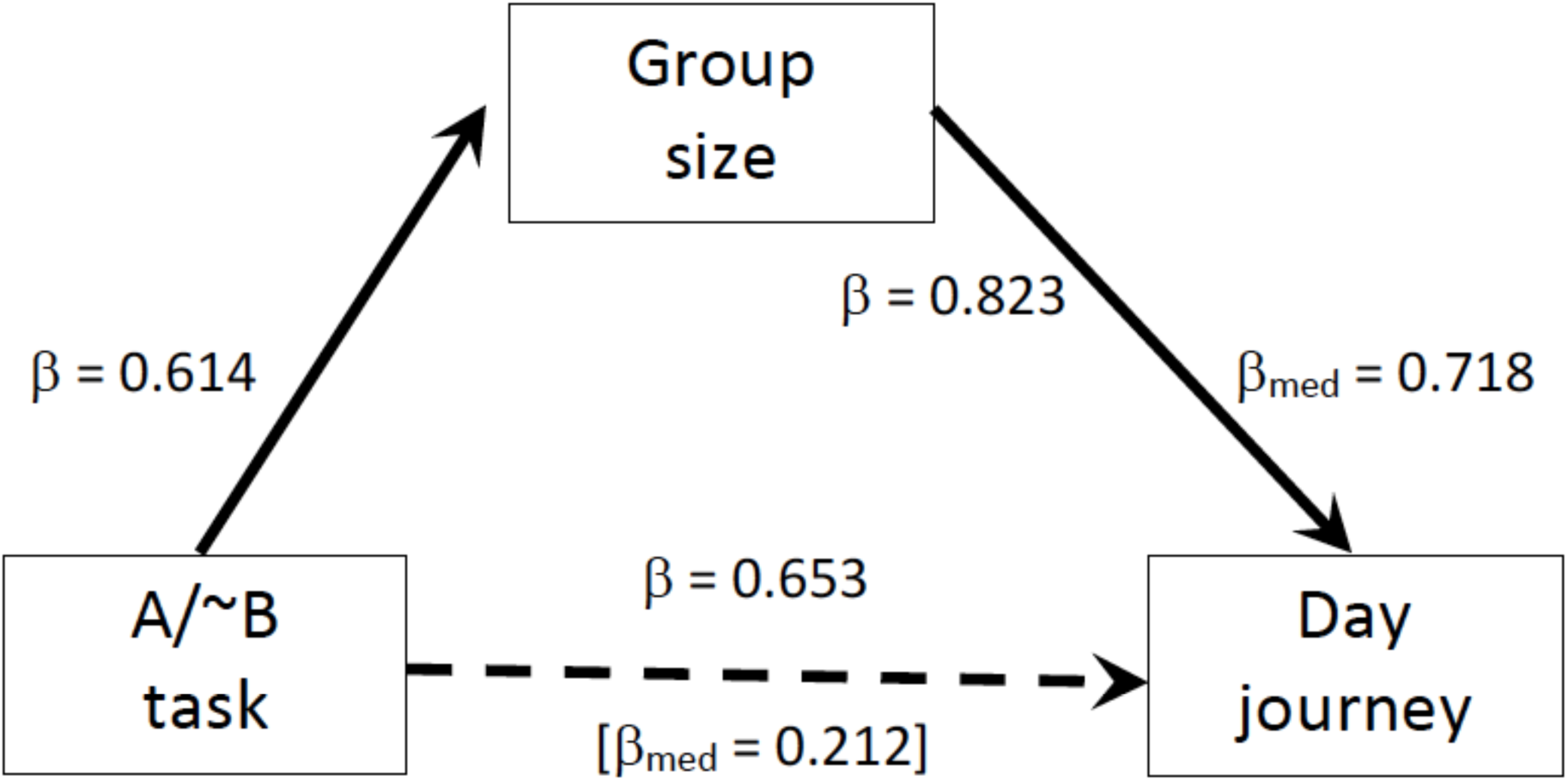
Mediation analysis of the relationship between the three variables in the social cluster for the A-not-B task in Table 4. A model with day journey length as the dependent variable gives a significantly better fit than one with group size as the dependent variable. βs are standardised slopes; β_med_ gives the standardised slope from the multiple regression equation with group size as the mediating variable.

Finally, we test the Passingham-Wise Conjecture using the two MacLean et al. [22] tasks for their full range of 36 mammal and bird species. Fig. 3a plots performance on the A-not-B task for the major taxonomic groupings in their full dataset. Performance varies significantly across mammalian orders (F_6,19_=3.73, p=0.013). It requires no statistical analysis to conclude that, just as Passingham & Wise (2011) suggested, self-control is unique to anthropoid primates: they are the only taxon whose scores lie significantly above chance (the dashed line at 33%). None of the non-anthropoid taxa (birds, rodents, carnivores, elephant and prosimian primates) perform at better than chance. Indeed, MacLean et al. (2014) themselves confirm this: they report that the correlation with brain size is not significant in the non-anthropoids (phylogenetically controlled regression, p=0.71), whereas there is a significant (p<0.01) correlation between brain size and self-control in the anthropoids (see also Dunbar & Shultz 2021a). Figure 3b plots the equivalent data for the cylinder task. In stark contrast to the A-not-B task, performance on the cylinder task does not differ significantly across the major taxonomic groups (F_5,25_=2.22, p=0.084). If anything, carnivores out-perform anthropoids on this task (albeit not significantly), with rodents running them a close second. Note that all three of these orders out-perform prosimian primates and birds. In short, this task is not indexing a cognitive skill that is specific to the primates.

**Fig. 3.**
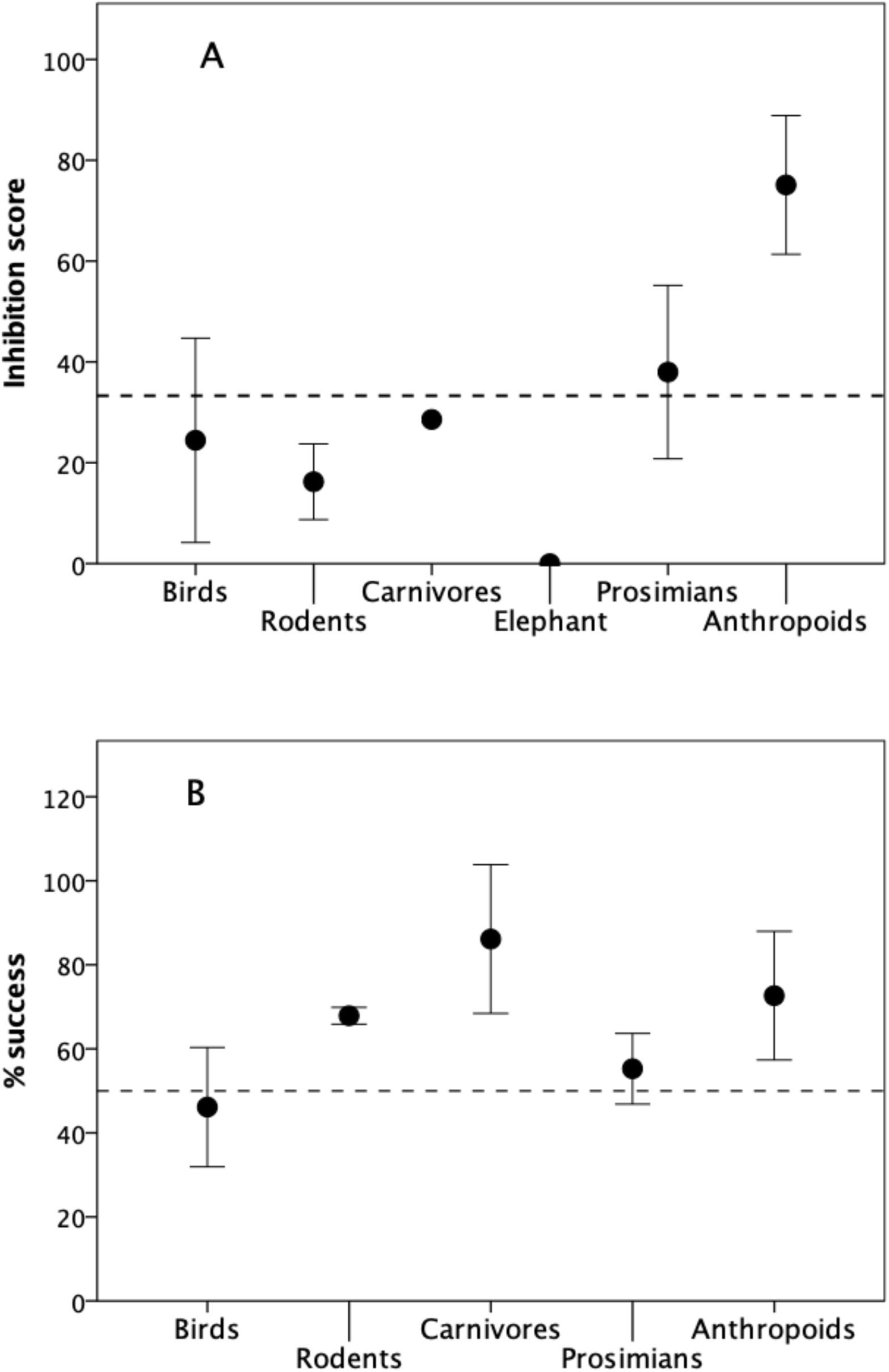
Performance on two cognitive tasks for different taxonomic groups for the two tasks in the MacLean et al. sample. (a) Mean (±2se) percentage success on the A-not-B inhibition task. The dashed horizontal line denotes the chance response rate at 33% (for a task in which the animal chooses between three locations). (b) Mean (±2 se) percentage success on the cylinder task. The dashed horizontal line denotes the chance response rate at 50% (for a task in which the animal chooses between one of two locations). Number of species sampled for each taxon: birds (7), rodents (2), carnivores (3), elephant (1), prosimians (8), anthropoid primates (15). Data from MacLean et al. (2014).

## Discussion

We have shown, using several very different indices of self-control from three independent databases, that inhibition (self-control) is closely correlated with two key variables that affect group coordination (group size and day journey length), but not with either of two indices that explicitly impact on food-finding decisions (percentage of fruit in the diet and home range size, with the latter being a proxy for decisions on where to find productive food patches). This suggests that the capacity to inhibit prepotent responses has less to do with food-finding *per se* (even if it might be secondarily beneficial once the capacity has evolved) than with the demands of maintaining group coordination whilst foraging. This concurs with human evidence that the ability to inhibit or delay gratification strongly predicts social skills, with poor capacity to do so being directly related to disruptive anti-social behaviour and an inability to maintain stable relationships (Tremblay et al. 1994; Moffet et al. 2001; Bari & Robbins 2012; Pearce et al. 2019), both of which are indicative of an inability to negotiate compromises. Although Jelbert et al. (2016) claimed that corvids can pass the A-not-B test, they in fact only did so when allowed to notice hand cues during reward switching; even then, only one out of eight birds actually managed this. In fact, five failed to pass the training phase, and, of the three that did pass the qualifying task, two responded only at chance levels on the test phase: this hardly seems to imply unqualified success. The issue here is likely the fact that anthropoid primates can use one-trial learning to infer general rules, whereas other taxa have to rely on much slower forms of associative learning (Passingham & Wise 2011).

In primates, including humans, both one-trial learning and inhibition are associated with the frontal pole (Briodman areas BA9/10: Passingham & Wise 2011; Passingham 2021) and the adjacent inferior prefrontal cortex (Aron et al. 2004), the part of the brain that has expanded most during the course of primate and human evolution – and is most dependent for full adult functionality on experience and learning during socialisation (Joffe 1997). More importantly, perhaps, neuropsychological studies of both primates and humans suggest that foraging-type decisions do not involve either inhibition or the frontal pole, but rather depend on the phylogenetically older anterior cingulate cortex further back in the brain (Brodman areas BA24 and 32-33: Kennerley et al. 2006; Holyroyd et al. 2009).

Tasks that primarily involve causal reasoning (such as the cylinder task and other detour tasks) give more ambiguous results, but mainly load with diet. The lack of any clear difference in performance on this task between mammals and birds reinforces the suggestion that these kinds of task index a generalised cognitive skill whose primary function is related to food-finding and other non-social forms of causality rather than specifically to social decision-making – or behavioural inhibition. It is noteworthy that birds and prosimians performed less well on the cylinder task than rodents, carnivores and anthropoid primates (Fig. 3b). The latter three orders probably engage in far more manipulation (or behavioural prediction, in the case of carnivores) of their food than the former. Although several authors have claimed that birds perform just as well as apes and monkeys on this task (Kabadayi et al. 2016, 2017; van Horik et al. 2018; Isaksson et al. 2018), the species that they tested (parrots, corvids, passerines) are all ones with large brains (for birds), possess sophisticated cognitive abilities and regularly manipulate food items to extract them from a matrix. Nonetheless, these results add weight to the suggestion that inhibition and causal reasoning have evolved independently of each other, and represent a case of mosaic evolution in cognitive skills and their underlying neural bases. The mediation analysis makes it clear that by far the best model of the causal relationships between the three variables in the social cluster is that the capacity to inhibit prepotent actions determines group size, and group size then determines day journey length. Biologically, this makes more sense than any alternative model. In the absence of the capacity to maintain group coherence, foraging groups will fragment and disperse, resulting in a reduction in the length of day journeys. This suggests that self-control plays a crucial role in managing group size and its constituent social relationships rather than influencing day journey length directly. The fact that most psychological studies invariably use food as a reward because this simplifies experimental designs (and allows experiments to be kept short) may be misleading in respect of the capacity’s function. More generally, the results suggest that greater caution needs to be exercised in making assumptions about what particular tests actually mean in terms of underlying cognitive skills.

These results feed into the longstanding distinction drawn between species that have stable social groups (congregations) and those that live in unstable herds (aggregations, or fission-fusion social systems). The former are characterized by intense affiliative relationships between individuals (mediated in primates by social grooming: Lehmann et al. 2007) and the constant monitoring of social partners (Dunbar & Shultz 2010). The capacity to inhibit and modulate behaviour is crucial for bonded relationships in that it ensures that the individuals synchronise their movements and hence do not lose contact with each other while foraging: when one stops to rest, the other must inhibit its desire to continue foraging and go to rest as well. Of the non-anthropoid species studied by MacLean et al. (2014), only elephants have bonded sociality above the level of monogamous pairbonds; however, elephants have a fission-fusion social system that does not depend on maintaining cohesion in large social groups (Moss et al. 2011), which may explain why they score very poorly on the inhibition task (Fig. 3a).

Maintaining the temporal coherence of bonded groups will not depend solely on the ability to inhibit prepotent responses. It is likely to involve other specialised forms of cognition, the most important of which will be understanding others’ intentions. In some Old World monkeys, for example, individuals make explicit bids, or suggestions, about the direction of group travel (often signalled by characteristic behaviours), with other group members then ‘voting’ on their preferences in order to arrive at a consensus (Stoltz & Saayman 1970; Sigg & Stolba 1981; Sueur & Pettit 2010; Sueur et al. 2011; Harel et al. 2021). Rapid assessment of another individual’s intentions will be essential not just to know how best to respond to their actions (e.g. to differentiate between threat and affiliation) but also to know whether a grooming partner intends to rest or continue moving. The capacity to infer the intentions of another individual is dependent on mentalising, a cognitive skill that is also confined to the anthropoid primates (Devaine et al. 2017). Like self-control, mentalising is neurophysiologically demanding (Dàvid-Barrett & Dunbar, 2013; Lewis et al. 2018) and dependent on specialised neural circuitry (Carrington & Bailey 2009; van Overwalle 2009). In humans, mentalizing skills of this kind are correlated both with the size of an individual’s social network (Stiller & Dunbar 2007; Powell et al. 2012) and with the volume of what has turned out to be the brain’s default mode neural network (Mars et al. 2012; Smallwood et al. 2021;

Roumazeilles et al. 2021). The default mode network is a substantial brain connectome involving prefrontal, parietal and temporal brain units, plus the limbic system and cerebellum and their substantial white matter connections, that humans share with at least the cercopithecine monkeys (Sallet et al. 2013; Mars et al. 2012, 2016; Rushworth et al. 2013; Li et al. 2014; Roumazeilles et al. 2020). The volume of this network correlates with the size of social groups and/or personal social networks in both humans (Lewis et al. 2011; Powell et al. 2012; Kanai et al. 2012; Kwak et al. 2018; Noonan et al. 2018; Kiesow et al. 2020) and anthropoid primates (Sallet et al. 2011; Meguerditchian et al. 2020; Testard et al. 2022). That mentalising and self-control seem to co-occur taxonomically suggests that they form part of a highly specialised suite of cognitive abilities dedicated principally to the maintenance of bonded social groups (Dunbar & Shultz 2021a).

## Supporting information

Sopplementary Methods and Analyses

## Acknowledgments

SS’s research is supported by a Royal Society University Research Fellowship (UF160725).

## Ethics Statements

### Ethics

All the data analysed in this paper are publicly available third party data. No ethics issues arise from the use of these data.

## Data Availability

All data generated or analysed during this study are included in this published article [and its supplementary information files].

