## Supplementary material for "The Social Role of Self-Control": Sopplementary Methods and Analyses

**R.I.M. Dunbar** and **Susanne Shultz**

#### Electronic Supplementary Material

### Supplementary Data

**Table S1. Group size, day journey length and the fissioning index for a sample of baboon study sites**

| Site | Group size | Day journey (km) | Fissioning index† | Source |
|---|---|---|---|---|
| *Awash Station, Ethiopia | 83.0 | 6.5 | 2 | Nagel 1973 |
| *ErerGota, Ethiopia | 83.0 | 8.9 | 2 | Sigg & Stolba 1981 |
| *Awash Filoha, Ethiopia | 190.0 | 7.5 | 2 | Swedell 2001 |
| Mt Assirik, Senegal | 247.0 | 7.9 | 2 | Byrne 1981; Sharman 1982 |
| Siminti, Senegal | 70.8 | 4.0 | 2 | Zinner et al. 2021 |
| Gashaka NP, Nigeria | 28.4 | 2.4 | 0 | Sommer & Ross 2010 |
| Metahara, Ethiopia | 87.0 | 5.8 | 1 | Aldrich-Blake et al. 1971 |
| Bole Valley, Ethiopia | 19.0 | 1.2 | 0 | R. Dunbar (unpublished) |
| Mulu, Ethiopia | 22.0 | 1.1 | 0 | R. Dunbar (unpublished) |
| Awash Falls, Ethiopia | 71.0 | 5.3 | 1 | Nagel 1973 |
| Budongo Forest, Uganda | 37.5 | 3.8 | 0 | Paterson 1976; pers. comm. |
| Chololo, Kenya ‡ | 102.0 | 5.6 | 1 | Barton 1990 |
| Gilgil, Kenya ‡ | 49.0 | 4.3 | 1 | Harding 1976 |
| Chololo STT 1986, Kenya | 102.0 | 5.6 | 1 | Barton 1990 |
| Chololo PHG 1995, Kenya | 25.0 | 4.6 | 0 | Kenyatta 1995 |
| Gombe NP, Tanzania | 43.0 | 2.4 | 0 | J. Oliver (pers. comm.) |
| Amboseli NP, Kenya [Hook] | 46.5 | 6.1 | 1 | D. Post (pers.comm.) |
| Ruaha NP, Tanzania | 72.0 | 6.8 | 1 | Collins 1984 |
| Mikumi NP (1995), Tanzania | 18.0 | 3.9 | 0 | Hawkins 1999 |
| Giants Castle, S. Africa | 11.8 | 0.9 | 0 | Henzi et al. 1992; R. Byrne (pers. comm.) |
| Cape Point, S. Africa | 85.0 | 7.9 | 2 | Davidge 1978 |
| Honnet, S. Africa | 77.0 | 9.0 | 2 | Stoltz & Saayman 1970 |
| Suikerbosrand, S. Africa | 78.0 | 4.1 | 2 | Anderson 1981 |
| R. Kuiseb, Namibia | 15.5 | 4.1 | 0 | Brain 1990 |
| Tsaobis, Namibia | 34.3 | 5.5 | 0 | King et al. 2008; G. Cowlishaw (pers. comm.) |

* *Papio hamadryas* † see Table S2
‡ based on a comparison of group size and the mean number of individuals within 10m of a focal adult

**Table S2. Fissioning index**

| Index | Definition |
|---|---|
| 0 | Group relatively compact during foraging, and always sleeps together; group spread during foraging always <200m |
| 1 | Group becomes dispersed during foraging (mean spread >200m), but always sleeps together |
| 2 | Group fragments during foraging, with sub-groups moving independently, with subgroups sometimes sleeping at separate sites |

For each study site, index is based on descriptions of foraging patterns given by primary sources

#### Diet data

We sourced our data on diet from Powell et al. (2017) rather than DeCasien et al. (2017) because we considered the Powell data compilation more reliable. However, the value of 8.5% that they give for *Macaca mulatta* is based on just one idiosyncratic high altitude study site. Other more typical habitats give much higher values of frugivory for this species. We searched for diet data for this species on GoogleScholar and located a further five studies (Table S3). We use the mean of these studies in our analyses.

**Table S3. Diet data for *Macaca mulatta***

| Study site | Country | % fruit in diet | Source |
|---|---|---|---|
| Taihangshan Reserve | China | 36.5 | Cui et al. (2018) |
| Nonggang Reserve | China | 30.0 | Tang et al. (2016) |
| *Murree Hills | Pakistan | 8.5 | Goldstein & Richard (1989) |
| Buxta Tiger Reserve | Bangladesh | 74.9 | Sengupta & Radhakrisha (2015) |
| Asola-Bhatti Sanctuary | India | 5.7 | Ganguly & Singh Chauhan (2018) |
| Siwalik Hills | India | 63.0 | Lindburg (1977) |
| **Mean** | | **37.2%** | |

* Site on which Powell et al. (2014) based their estimate.

Powell et al. (2014) do not give a value for percent of fruit in diet for *Saguinus oedipus*. We use the value given for this species by Garber (1984).

We did not use Powell et al. (2017) as a source of data for day journey length or home range size because, although their values for day journey correlate significantly with those we compiled from Smuts et al. (1987) and Campbell et al. (2008) ($r=0.887$, $p=0.003$), those for range size, in particular, appear to be based on a very selective sampling of study sites. Our sources are based on a wider range of primary sources, and are likely to be more representative.

**Exclusions and Anomalous Group Sizes**

*Daubentonia* was omitted from the MacLean et al. dataset because of doubts over the correct group size to use. Although *Daubentonia* (a very rare and difficult to study nocturnal prosimian) has been consistently listed with a group size of N=1 in most comparative datasets because it forages solitarily, in fact field studies have suggested that it actually lives in much larger communities (neighbourhoods) as do all the other nocturnal prosimians (Iwano, 1991; Ancrenaz, Lackman-Ancrenaz & Mundy, 1994; Sterling & McCreless, 2006). However, there are no reliable estimates of what this group size actually is (though values around N=8 have been suggested – i.e. considerably larger than the conventionally cited value of N=1).

*Papio hamadryas* was omitted from the MacLean et al. dataset for related reasons. This species is unusual for a baboon in that it lives in a multilevel society with at least four different grouping levels (Hill et al. 2008). It has never been clearly determined which of these grouping levels is the correct one to use in comparatrive studies. Although the band (mean size N=84.5) is commonly cited, there are good cognitive and socio-demographic grounds for considering the clan (mean size N=24.0) as the more appropriate natural grouping (Dunbar & Shultz 2021a). It is important to note that mean band size is 3.3 SDs above the overall mean for the MacLean sample and the species' mean day journey length is 3.5 SDs above the sample mean. It is normally customary to exclude values that lie >2 SDs from the mean. For present purposes, we excluded the species, but give results for separate analyses with the two candidate group sizes in the ESM.

The orang utan (*Pongo*) provides another potentially problematic case: most comparative databases give a group size of N=1 for this species on the grounds that, like *Daubentonia*, it typically forages alone. However, these genus currently occupies a retreat habitat on the limits of its biogeographical range (see Dunbar 1988; Calne et al. 2010). Subfossil orangs on the Chinese mainland occupied woodland rather than forest habitats and were likely much more terrestrial than they are now (Smith & Pilbeam 1980; Harrison et al. 2002) and so almost certainly foraged in much larger groups. In fact, this species is more intensely social than the gorilla and is commonly kept in groups of up to 7 animals in captivity (Edwards & Snowdon 1980; Lardeux-Gilloux 1997). Indeed, some contemporary populations forage in larger groups (Galdikas 1985; Delgado & van Schaik 2000; Setia et al. 2009). Recent studies provide compelling evidence that communities of 12-15 are typical (Mackinnon 1974; Singleton & van Schaik 2001), and this value fits extremely closely with the size we would predict given its neocortex size and the ape social brain relationship (Dunbar et al. 2018; Dunbar & Shultz 2021a). As with *Daubentonia*, using the conventional group size of N=1 risks confounding foraging group size with social group size (Dunbar & Shultz 2023). We use the value of N=14 given by Dunbar et al. (2018).

### Supplementary analyses

#### Bivariate correlations

**Table S4. Bivariate Pearson correlations between the cognitive tasks provided by Amici et al. (2008, 2018) and MacLean et al. (2014)**

| | Amici et al. (2008) | Amici et al. (2018) | MacLean et al. (2014) | |
|---|---|---|---|---|
| | Causal reasoning | Social inhibition | A-not-B | Cylinder |
| Amici Inhibition (mean) | 0.454 | 0.577 | 0.385 | 0.661 |
| p | (0.153†) | (0.115‡) | (0.197†) | (0.165*) |
| Causal reasoning (mean) | | **0.833** | 0.581 | -0.560 |
| p | | **(0.020‡)** | (0.086†) | (0.780†) |
| Social inhibition | | | 0.110 | -0.028 |
| p | | | (0.418‡) | (0.986*) |
| A-not-B | | | | -0.744 |
| p | | | | (0.628*) |

All p-values are the probability of a positive (1-tailed) correlation; bold values are significant.
† N=7; ‡ N=6; * N=4

**Table S5. Bivariate Pearson correlations for the Stevens (2014) and MacLean et al. (2014) tasks**

| | Diet | Group size | Day journey | Home range |
|---|---|---|---|---|
| Go/no-go | r=0.206<br>p=0.544<br>N=11 | r=0.676<br>p=0.022<br>N=11 | r=0.866<br>p=0.001<br>N=11 | r=0.258<br>p=0.444<br>N=11 |
| A-not-B | r=-0.072<br>p=0.758<br>N=21 | r=0.637<br>p=0.002<br>N=21 | r=0.603<br>p=0.004<br>N=21 | r=0.461<br>p=0.041<br>N=20 |
| Cylinder | 0.428<br>p=0.076<br>N=18 | 0.519<br>p=0.027<br>N=18 | 0.512<br>p=0.030<br>N=18 | 0.443<br>p=0.066<br>N=18 |
| Diet | | -0.086<br>p=0.695<br>N=23 | -0.197<br>p=0.366<br>N=23 | -0.06<br>p=0.769<br>N=23 |
| Group size | | | 0.813<br>p<0.001<br>N=24 | 0.365<br>p=0.087<br>N=23 |
| Day journey | | | | 0.242<br>p=0.265<br>N=23 |

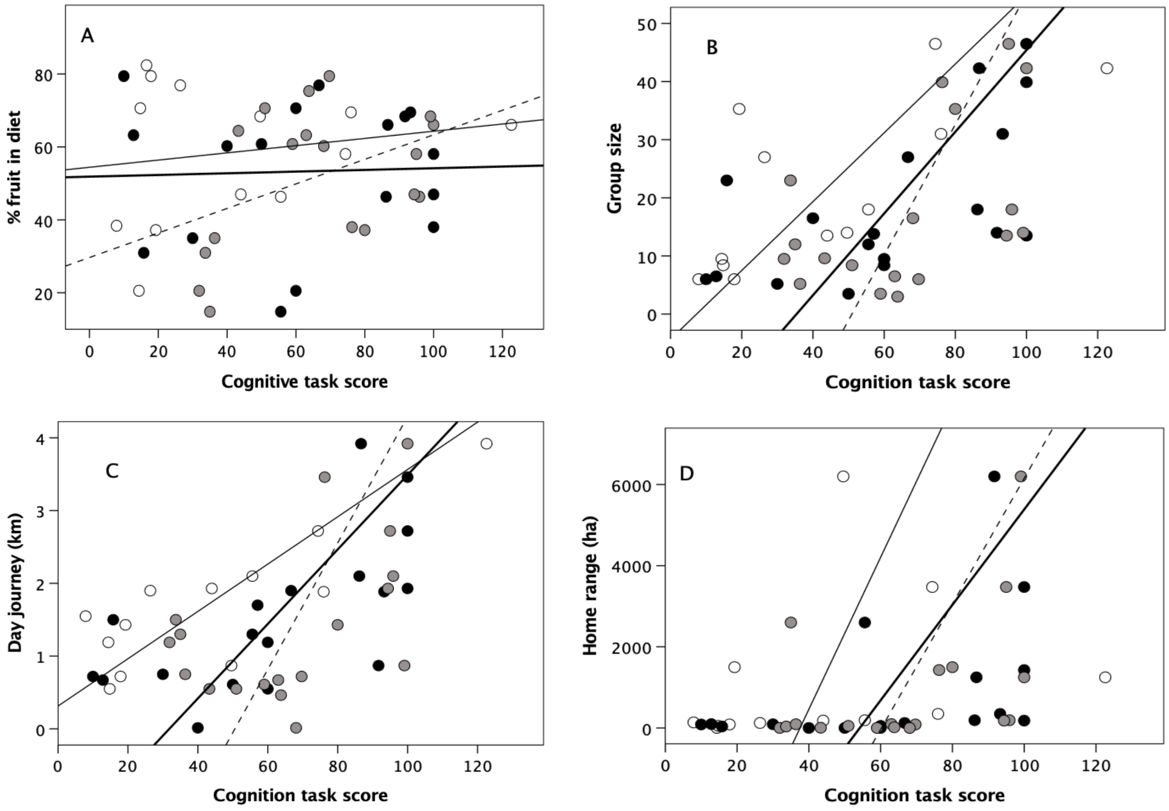


**Figure S1**. Cognition score as predictor of (a) diet (% fruit), (b) mean group size, (c) mean day journey length (km), and (d) mean home range area (ha) for individual species. Unfilled circles, thin solid line: MacLean et al. A-not-B task; grey circles, dashed line: MacLean et al. cylinder task; filled circles, thick line: Stevens Go/No-go task. Lines are OLS regressions.

**Table S6. PCA factor loadings (with varimax rotation) for the mean value of the social inhibition task from Amici et al. (2008)**

| Factors: | 1 | 2 |
|---|---|---|
| Inhibition tasks (mean) † | **0.891** | 0.048 |
| Group size | **0.921** | 0.128 |
| Day journey | **0.885** | -0.403 |
| Diet (% fruit) | -0.024 | **0.780** |
| Range size | -0.004 | **0.777** |
| Species sampled | 7 | |
| Variance explained (%) | 76.4 | |

Extraction based on λ = 1.0. Bold font indicates variables that have a strong positive load on the same factor.

**Table S7a. PCA factor loadings (with varimax rotation) for the variables for each of the three datasets, with orang utan (*Pongo*) excluded.**

| | Stevens (2014) Go/No-go task | | MacLean et al. (2014) A-not-B task only | | Amici et al. (2008) Inhibition task ‡ | |
|---|---|---|---|---|---|---|
| Factors: | 1 | 2 | 1 | 2 | 1 | 2 |
| Inhibition task | **0.889** | 0.318 | **0.831** | 0.009 | **0.864** | -0.156 |
| Group size | **0.934** | -0.165 | **0.933** | 0.106 | **0.933** | 0.322 |
| Day journey | **0.935** | 0.150 | **0.923** | -0.022 | **0.859** | -0.257 |
| Diet (% fruit) | 0.054 | **0.928** | 0.035 | **0.967** | 0.067 | **0.982** |
| Range size | **0.748** | -0.445 | 0.698 | -0.415 | **0.797** | -0.014 |
| Species sampled | 12 | | 17 | | 6 | |
| Variance explained | 86.2% | | 80.5% | | 83.1% | |

Extraction based on λ = 1.0. Bold font indicates variables that have a strong positive load (>0.700) on the same factor.
‡ mean of two inhibition tasks (A-not-B and delayed gratification tasks)

**Table S7b. PCA factor loadings (with varimax rotation) for the variables for the MacLean et al. dataset, with *Papio hamadryas* included at two different grouping levels.**

| | MacLean A-not-B task | | | |
|---|---|---|---|---|
| Grouping level: | Band (N=84.5) | | Clan (N=24.0) | |
| Factors: | 1 | 2 | 1 | 2 |
| Inhibition task | 0.568 | 0.535 | **0.830** | 0.077 |
| Group size | **0.928** | 0.059 | **0.852** | 0.050 |
| Day journey | **0.947** | -0.056 | **0.735** | -0.310 |
| Diet (% fruit) | -0.275 | **0.728** | 0.040 | **0.969** |
| Range size | 0.414 | 0.591 | 0.603 | 0.075 |
| Species sampled | 22 | | 22 | |
| Variance explained | 70.0% | | 67.4% | |

Extraction based on λ = 1.0. Bold font indicates variables that have a strong positive load (>0.700) on the same factor.

**Genus level analysis**

Although there is negligible phylogenetic signal in any of the data and analyses of the data with and without phylogenetic control yield identical results (MacLean et al. 2014; Stevens 2014), we nonetheless further checked whether phylogenetic inertia might distort our results by re-analysing the data as genus-level averages. In fact, there are only three genera with more than a single species sampled in either of the two datasets. The results are given in Table S7. As before, a factor analysis with λ=1 explains 70-74% of the variance, with the Go/No-Go and A-not-B tasks loading with group size and day journey length as before, and the cylinder task loading with diet and home range size. The only difference is that range size loads more strongly with diet on factor 2 in both datasets than was the case with the larger sample.

**Table S8. PCA factor loadings (with varimax rotation and λ>1) for the five variables for each of the two datasets for mean genus-level data. Bold font indicates variables that load together on the same factor.**

| | Go/No-go task | | A-not-B task | | Cylinder task | |
|---|---|---|---|---|---|---|
| Factors: | 1 | 2 | 1 | 2 | 1 | 2 |
| Cognitive task | **0.774** | 0.583 | **0.853** | 0.236 | 0.225 | **0.902** |
| Diet | -0.252 | **0.893** | -0.224 | **0.801** | -0.648 | **0.666** |
| Group size | **0.865** | -0.170 | **0.783** | -0.333 | **0.880** | 0.216 |
| Day journey | **0.881** | 0.106 | **0.853** | -0.188 | **0.805** | 0.277 |
| Range size | 0.200 | **0.562** | **0.517** | **0.543** | 0.231 | **0.581** |
| Variance explained | 74.4% | | 70.5% | | 73.3% | |
